# Targeting mitochondrial dysfunction using methylene blue or mitoquinone to improve skeletal aging

**DOI:** 10.1101/2023.06.16.545146

**Authors:** Sher Bahadur Poudel, Dorra Frikha-Benayed, Ryan R. Ruff, Gozde Yildirim, Manisha Dixit, Ron Korstanje, Laura Robinson, Richard A. Miller, David E. Harrison, John R. Strong, Mitchell B. Schaffler, Shoshana Yakar

## Abstract

Declines in mitochondrial content and impaired cytochrome c oxidase activity (complex IV) can result in reduced energy metabolism and increased levels of oxidants. Methylene blue (MB) is a well-established antioxidant that has been shown to improve mitochondrial function in both in vitro and in vivo settings. Mitoquinone (MitoQ) is a selective antioxidant that specifically targets mitochondria and effectively reduces the accumulation of reactive oxygen species (ROS) within cells.

To investigate the effect of acute and long-term administration of MB on skeletal morphology, we administered MB to aged (18 months old) female C57BL/J6 mice, as well as to adult male and female mice with a genetically diverse background (UM-HET3). Additionally, we used MitoQ as an alternative approach to target mitochondrial oxidative stress during aging in adult female and male UM-HET3 mice.

Although we observed some beneficial effects of MB and MitoQ in vitro, the administration of these compounds in vivo did not alter the progression of age-induced bone loss. Specifically, treating 18-month-old female mice with MB for 6 or 12 months did not have an effect on age-related bone loss. Similarly, long-term treatment with MB from 7 to 22 months or with MitoQ from 4 to 22 months of age did not affect the morphology of cortical bone at the mid-diaphysis of the femur, trabecular bone at the distal-metaphysis of the femur, or trabecular bone at the lumbar vertebra-5 (L5) in UM-HET3 mice.

Based on our findings, it appears that long-term treatment with MB or MitoQ alone, as a means to reduce skeletal oxidative stress, is insufficient to inhibit age-associated bone loss. This supports the notion that interventions solely with antioxidants may not provide adequate protection against skeletal aging.

## Introduction

The existing literature supports the notion that oxidative stress and mitochondrial dysfunction are contributors to impaired skeletal aging. This concept is affirmed by randomized clinical studies (summarized in^1^), which have shown a correlation between consumption of an antioxidant-rich diet and risk of osteoporosis^2,3^. Furthermore, a study of postmenopausal women with osteoporosis revealed decreased activity of the antioxidant enzyme glutathione reductase in serum in comparison to non-osteoporotic postmenopausal women, as well as increased plasma levels of the oxidative stress marker, malondialdehyde^4^. In addition, a range of in vitro and in vivo animal studies (reviewed in ^5,6^) have revealed a negative association between oxidative stress and bone mineral density (BMD). In vitro studies show that H□O□-induced oxidative stress adversely affected the proliferation, differentiation, and mineralization of rat primary osteoblast cultures, yet the effects were partially alleviated by supplementation with the antioxidant amino acid derivative, N-acetyl cysteine (NAC)^7^. Overall, it appears that disruptions in redox homeostasis have a detrimental impact on bone cell differentiation, function, and overall skeletal health. As a result, there is a growing interest in exploring the potential prophylactic use of antioxidants as a means to prevent osteoporosis and age-related bone loss.

Methylene blue (MB) is an FDA-approved medication that has been widely used in clinical practice for over a century. Recent research has revealed that MB acts directly on mitochondria^8^. MB has a low redox potential, and in the presence of molecular oxygen, the reduced form of MB (MBH_2_) bypasses the production of harmful superoxide radicals. Instead, it enhances the efficiency of the Electron Transport Chain (ETC) by increasing the activity of mitochondrial complex IV. This leads to improved mitochondrial function and helps maintain the integrity of these crucial cellular structures (**Figure 1A**).

**Figure 1:**
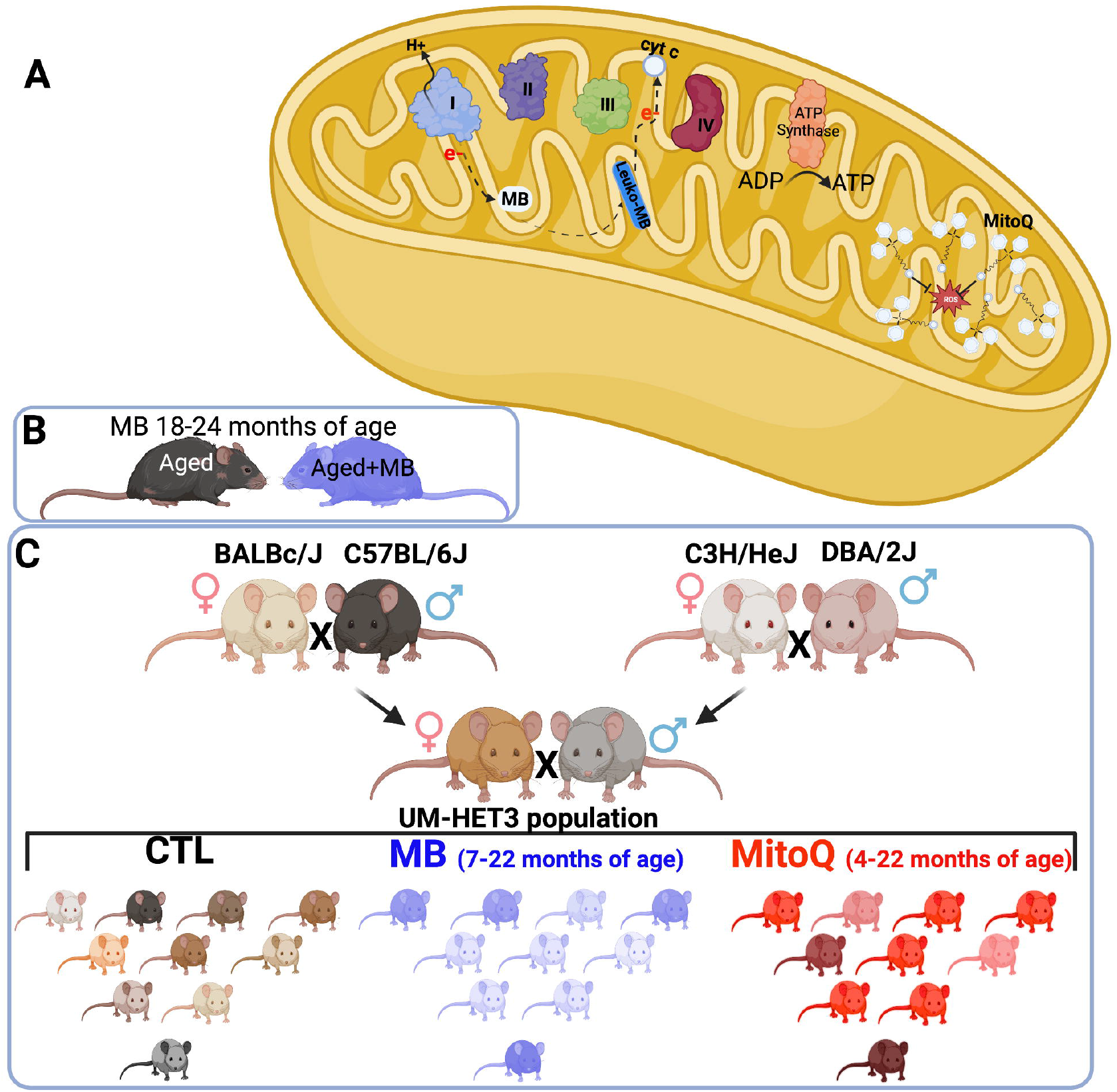
Experimental design. (A) Schematic illustration of MB and MitoQ actions in mitochondria. Experimental design of studies both inbred (C57BL/6J) (B) and outbred (UM-HET3) (C) mouse lines.

Previous studies have demonstrated various beneficial effects of MB. MB led to significant increase in “maximal lifespan” (proportion of mice alive at the 90th percentile) of genetically diverse UM-HET3 female mice^9^. It has been shown to improve memory function in aging individuals^10-13^ and has the potential to counteract cognitive impairments associated with conditions such as Alzheimer’s disease. MB can also inhibit the production of amyloid-beta, a protein implicated in Alzheimer’s pathology, and rescue cognitive defects^14,15^. Additionally, MB has been found to promote wound healing^16^, stimulate the proliferation of skin cells, and reduce markers of aging in the skin^17,18^. Notably, MB has shown promise in the treatment of Hutchinson-Gilford Progeria Syndrome (HGPS), a genetic disorder that is in some respects a model of accelerated aging^19^.

Mitoquinone (MitoQ) is an antioxidant that targets mitochondria. It is a lipophilic conjugated compound that has the ability to accumulate in high concentrations within the mitochondria (**Figure 1A**). Once inside the mitochondria, MitoQ activates ubiquinone oxidoreductases, which are enzymes involved in the electron transport chain. This activation leads to the reduction of ubiquinone to its active antioxidant form called ubiquinol^20^.

The administration of MitoQ did not result in a significant effect on lifespan when tested in UM-HET3 mice^9^; however, evidence from several studies suggests that MitoQ is beneficial for bone health. In db/db mice and in mice fed a high-fat diet, MitoQ was found to inhibit bone loss. Additionally, in vitro studies using bone marrow stromal cells from db/db mice showed that MitoQ significantly improved osteogenic differentiation. This improvement was associated with decreases in the levels of ROS and other markers of oxidative stress^21^. MitoQ was also able to attenuate osteoblast apoptosis caused by advanced glycation end products (AGEs)^22^. Moreover, MitoQ was shown to prevent increased oxidative stress and stimulate mitophagy in a mouse model of periodontitis, both of which are beneficial to bone formation and remodeling^23^. In vitro and in vivo studies also revealed that MitoQ can effectively reduce mitochondrial oxidative damage caused by erastin-induced ferroptosis and myocardial injury respectively^24,25^.

In collaboration with The Jackson Laboratory Center for Aging Research, we conducted an experiment where we administered Methylene Blue (MB) to 18-month-old female mice of the C57BL6 strain. We then monitored changes in bone structure for a period of 6 or 12 months after the treatment (**Figure 1B**).

Additionally, we collaborated with the Interventions Testing Program (ITP), which is supported by the National Aging Institute (NIA), to investigate the effects of lifelong treatment with MB or MitoQ on skeletal structure in genetically diverse UM-HET3 mice (**Figure 1C**). To assess the skeletal phenotype, we analyzed the appendicular skeleton (specifically the femur) and the axial skeleton (specifically the lumbar vertebra-5) using micro-computed tomography (micro-CT).

## Results

### Effects of MB and MitoQ on mesenchymal stem cell (MSC) viability, in vitro osteogenesis, and mitochondrial respiration in osteoblasts

To investigate the effects of MB and MitoQ on various aspects of osteogenesis, we conducted in vitro experiments using mesenchymal stem cells (MSCs) and differentiated osteoblasts from femur and tibia of 7-month-old UM-HET3 mice. First, we assessed the viability of MSCs by culturing them with increasing concentrations of MB or MitoQ for 48 hours. Our results, shown in **Supplement Figure 1**, indicated that MSC viability was not significantly affected by doses of 0.125-0.5 µM of MB or MitoQ. Next, we examined the effect of MB and MitoQ on in vitro osteogenesis. MSCs were induced to differentiate into osteoblasts in the presence of varying concentrations of MB or MitoQ for 18-28 days. We evaluated the presence of alkaline phosphatase (Alk-Phos) positive colonies as an indicator of osteogenic activity at 18 days of culture. Importantly, no significant differences in Alk-Phos staining were observed between the control group and those treated with MB or MitoQ at concentrations ranging from 0.125-0.5 µM (**Figure 2A-C**). To investigate the effects of MB and MitoQ on osteoclastogenesis, we isolated bone marrow hematopoietic cells and stimulated them with RANKL and M-CSF in the presence of increasing concentrations of MB and MitoQ. Our findings demonstrated that MB and MitoQ inhibited in vitro osteoclast differentiation in a dose-dependent manner, as evidenced by a reduced number of multinucleated, TRAP positive cells at concentrations above 0.25 µM (**Figure 2D-F**). Additionally, we examined the effect of MB and MitoQ on the respiration of mature osteoblast cells. After 14 days in culture, mature osteoblasts were exposed to different concentrations of MB or MitoQ for 48 hours and then subjected to a mitochondrial stress assay using the Seahorse bioanalyzer. Interestingly, we observed that 0.5 µM of MB or MitoQ resulted in a reduction of basal oxygen consumption rate (OCR) (**Figure 2G-I**) and maximal respiration (**Figure 2J**) in differentiated osteoblasts, although these changes did not reach statistical significance.

**Figure 2:**
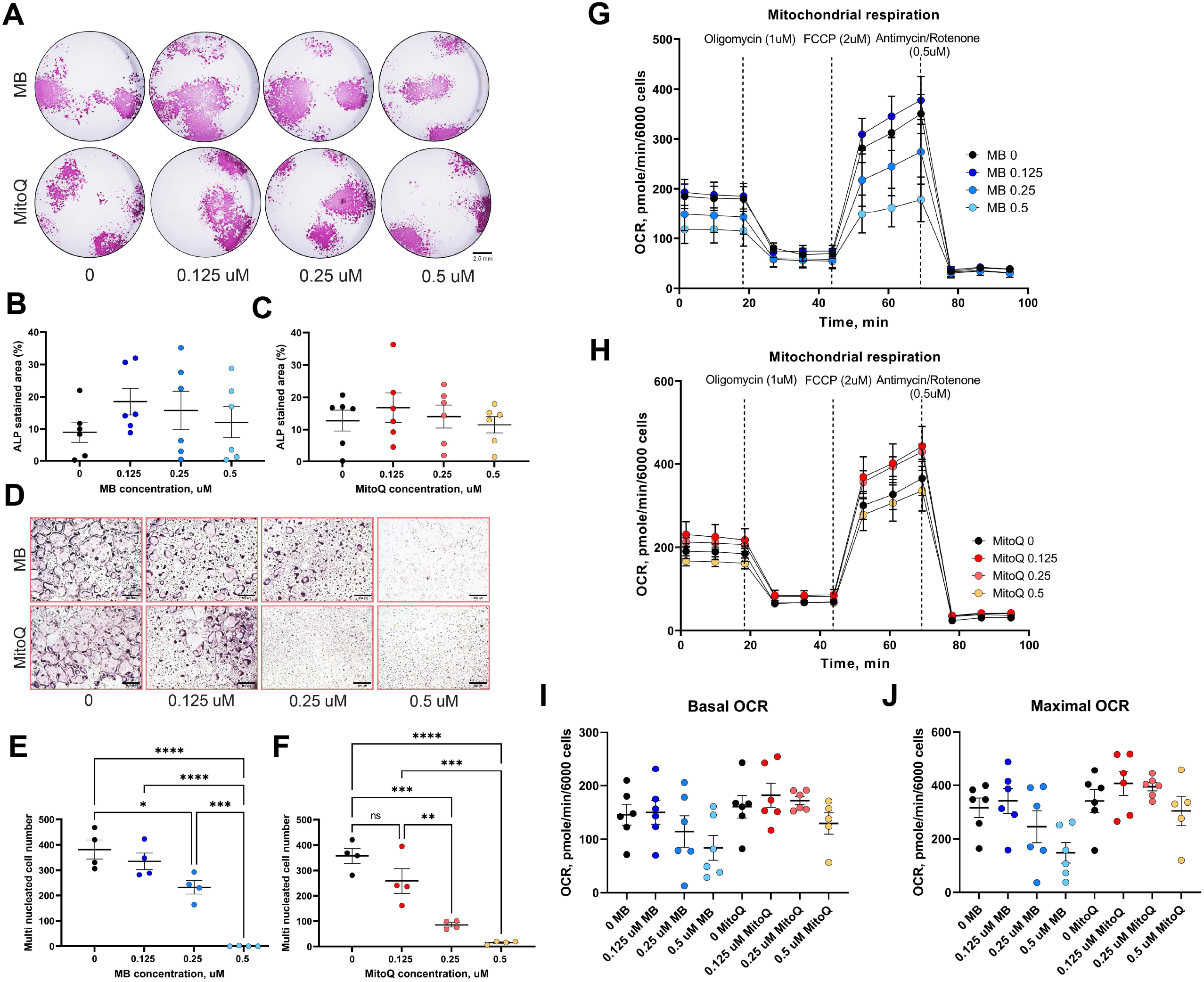
Effects of MB and MitoQ on osteogenesis in vitro. BM cells from 6-7 months old UM-HET3 mice were seeded in 24 well plate at a density of 1.5 10^6^ cells/well. Adherent cells were subjected for osteoblast differentiation treating with osteogenic factors in the presence or absence of MB or Mito Q at different concentrations (0, 0.125, 025 and 0.5 uM). (**A**) On day 18 in culture, cells were stained for Alkaline-phosphatase (Alk-Phos). The area of Alk-Phos positive colonies treated with (**B**) MB or (**C**) MitoQ were quantified in the wells were measured. (**D**) Non-adherent BM mononuclear cells were separated by Ficoll-Paque density gradient and seeded on 96 wells plate and induced for osteoclast differentiation with the supplementation of RANKL, M-CSF in the presence or absence of (**E**) MB or (**F**) MitoQ (0, 0.125, 025 and 0.5 uM). Multinucleated osteoclasts (>3 nuclei/cell) formed after 5 days in culture were visualized using TRAP staining kit and counted. Differentiated osteoblasts (14 days in culture) were treated with (**G**) MB or (**H**) MitoQ for 48 hours and oxygen consumption rate (OCR) was measured using mitochondrial stress assay kit. (**I**) Basal OCR and (**J**) maximal OCR were recorded along the assay. Data presented as mean+/-SEM. We used n=6 mice for assays in **A**-**C**, n=4 for assays in **D**-**F**, n=6 for mitochondrial stress assay in **G**-**J**. Data tested by multivariate ANOVA. Significance accepted at p<0.05 (*p<0.05, **p<0.01, ***p<0.001, ****p<0.0001).

In summary, our in vitro studies indicated that MB and MitoQ did not significantly affect MSC viability or in vitro osteogenesis at concentrations ranging from 0.125-0.5 µM. However, these compounds demonstrated dose-dependent inhibition of osteoclast differentiation. Furthermore, 0.5 µM of MB or MitoQ appeared to reduce basal OCR and maximal respiration in differentiated osteoblasts, although these changes were not statistically significant.

### Effects of MB on skeletal morphology of the axial and appendicular skeleton of aged female C57BL/6J mice

To investigate the effects of MB on bone morphology during aging, we conducted a study using 18-month-old female C57BL/6J mice. The mice were treated with 250 µM of MB, administered via drinking water for either 6 or 12 months. We dissected bones and tissues at 18, 24, or 30 months of age. Additionally, we included a group of young control females at 5 months of age to evaluate the changes from young to aged mice. The dissected bones were subjected to micro-CT to study bone morphology at three different sites: cortical bone at the femur mid-diaphysis, trabecular bone of the appendicular skeleton at the femur distal metaphysis, and trabecular bone of the axial skeleton at the lumbar vertebra-5 (L5).

Our findings revealed age-associated bone loss at all skeletal sites (**Figure 3**). In the control females, the total cross-sectional area (T.Ar) increased by 35%, bone area (B.Ar) increased by 24%, and marrow area (M.Ar) increased by 40% from 5 to 18 months of age (data not shown) (**Figure 3A, B**). This resulted in a 10% decrease in the B.Ar/T.Ar ratio between 5 and 18 months of age (**Figure 3D**). Cortical thickness (C.Th) remained similar between 5 and 18 months of age but decreased by 10% between 18 and 24 months and by 30% between 18 and 30 months of age (**Figure 3C**). Mice treated with MB showed similar changes to their corresponding controls at the same age.

**Figure 3:**
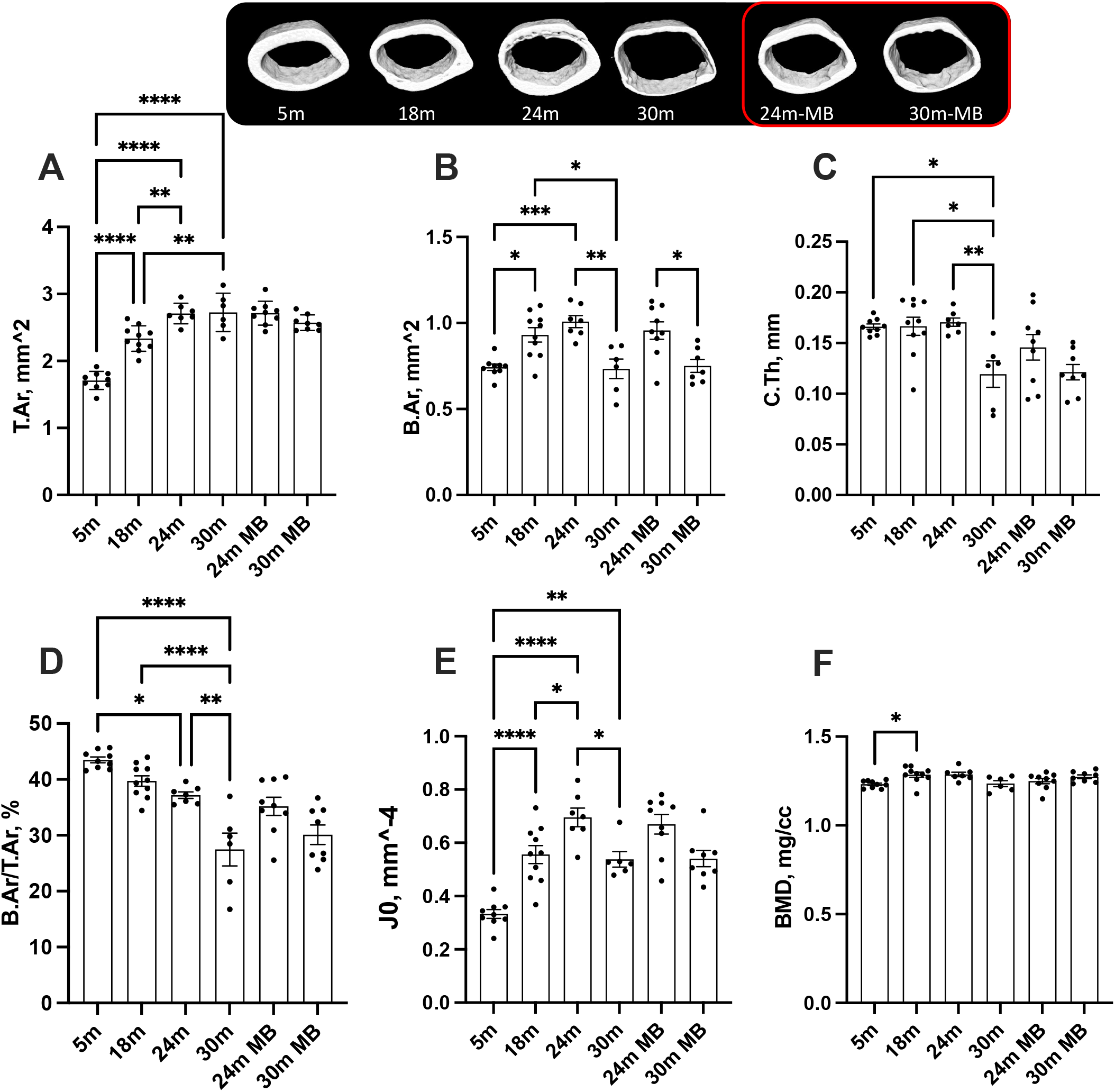
Administration of MB during aging does not alter cortical bone morphology of the appendicular skeleton. Eighteen-months old female C57BL/6J were divided into two groups. One group was exposed to regular drinking water and the other to 250 uM MB-containing water. Mice were sacrificed at basal (18 months), 24, or 30 months of age. An additional group of female mice at 5 moths of age, served as young controls. Bones were dissected and subjected to micro-CT. Femurs were scanned at 9.7 um resolution. Cortical bone parameters were evaluated at a 2mm^3 volume at the femoral mid-diaphysis including, (A) T.Ar-total cross-sectional area, (B) B.Ar-bone area, (C) C.Th-cortical bone thickness, (D) B.Ar/T.Ar-cortical bone volume/total volume, (E) J0-Polar moment of inertia, and (F) BMD-bone mineral density. Data presented as mean+/-SEM. 5 months-old females n=9, 18 months-old females n=10, 24 months-old females n=7, 30 months-old females n=6, 24 months-old MB-treated females n=9, 30 months-old MB-treated females n=8. Data tested by multivariate ANOVA presented as mean+/-SEM. Significance accepted at p<0.05 (*p<0.05, **p<0.01, ***p<0.001, ****p<0.0001).

At 5 months of age, control females exhibited low levels of trabecular bone volume/total volume (BV/TV) at the distal femur metaphysis, ranging between 5-6% (**Figure 4A**). Furthermore, we observed decreases in bone mineral density (BMD) in the trabecular bone compartment at the distal femur, along with decreased trabecular thickness (Tb.Th) with age, regardless of MB treatment (**Figure 4B, C**). However, trabecular number (Tb.N) did not show significant differences with age or treatment (**Figure 4D**). Similarly, the trabecular bone of the L-5 revealed a 35% reduction in BV/TV, 18% reduction in BMD, and a twofold decrease in Tb.N with age, irrespective of MB treatment (**Figure 5A-C**). Tb.Th did not exhibit significant differences between the age or treatment groups (**Figure 5D**). Overall, MB did not protect against age-induced bone loss.

**Figure 4:**
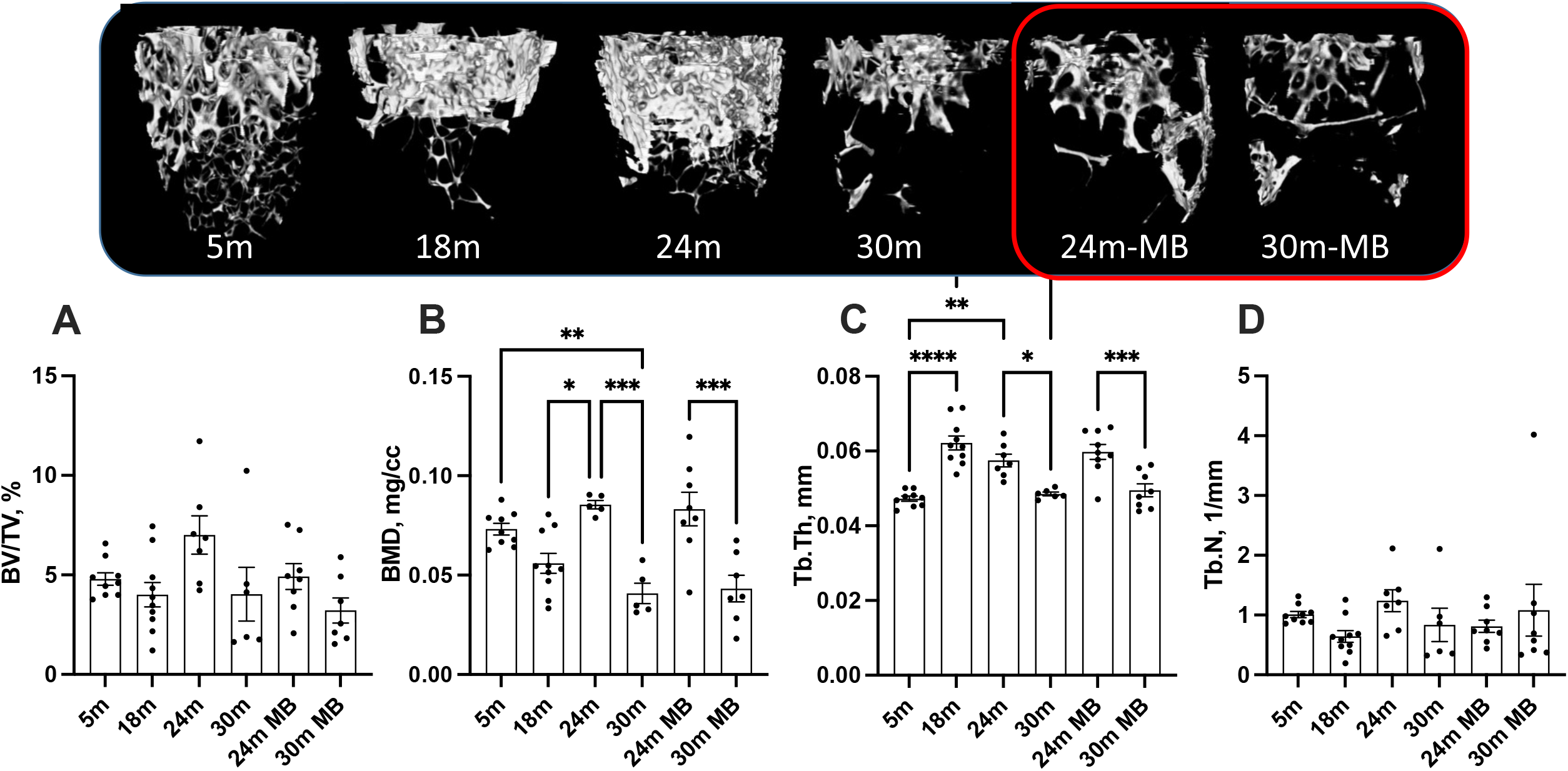
Administration of MB during aging does not alter trabecular bone morphology of the appendicular skeleton. Eighteen-months old female C57BL/6J were divided into two groups. One group was exposed to regular drinking water and the other to 250 uM MB-containing water. Mice were sacrificed at basal (18 months), 24, or 30 months of age. An additional group of female mice at 5 moths of age, served as young controls. Bones were dissected and subjected to micro-CT. Femurs were scanned at 9.7 um resolution. Trabecular bone parameters were taken at a 2mm^3 volume 200 um proximal to the femur distal metaphysis including (A) BV/TV-bone volume/total volume, (B) BMD-bone mineral density, (C) Tb.Th-trabecular thickness, and (D) Tb.N-trabecular number. Data presented as mean+/-SEM. 5 months-old females n=9, 18 months-old females n=10, 24 months-old females n=7, 30 months-old females n=6, 24 months-old MB-treated females n=8, 30 months-old MB-treated females n=7. Data tested by multivariate ANOVA presented as mean+/-SEM. Significance accepted at p<0.05 (*p<0.05, **p<0.01, ***p<0.001, ****p<0.0001).

**Figure 5:**
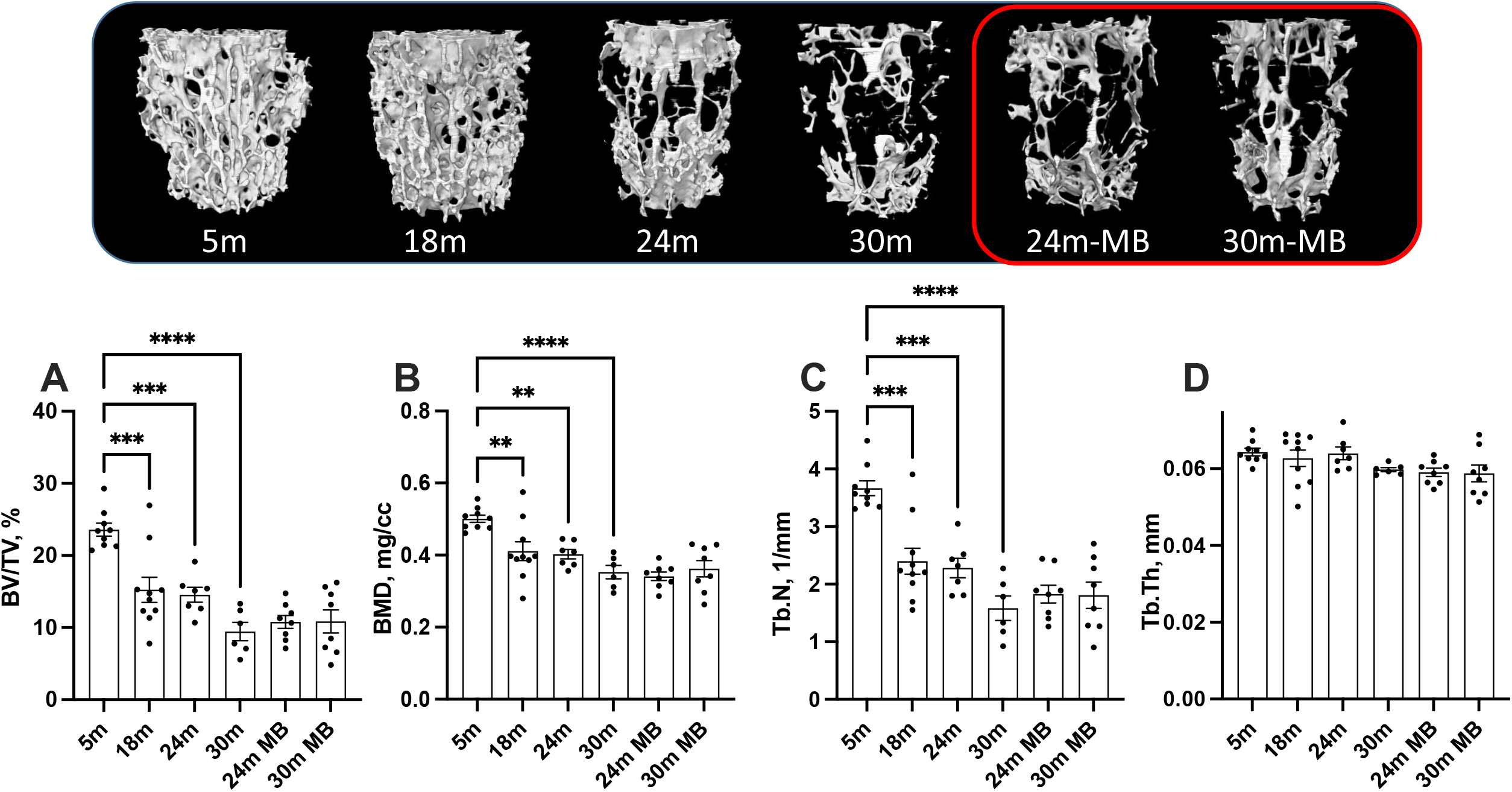
Administration of MB during aging does not alter trabecular bone morphology of the axial skeleton. Eighteen-months old female C57BL/6J were divided into two groups. One group was exposed to regular drinking water and the other to 250 uM MB-containing water. Mice were sacrificed at basal (18 months), 24, or 30 months of age. An additional group of female mice at 5 moths of age, served as young controls. Bones were dissected and subjected to micro-CT. Trabecular bone parameters of the L5 vertebra were taken at the vertebral body including, (A) BV/TV-bone volume/total volume, (B) BMD-bone mineral density, (C) Tb.Th-trabecular thickness, and (D) Tb.N-trabecular number. 5 months-old females n=9, 18 months-old females n=10, 24 months-old females n=7, 30 months-old females n=6, 24 months-old MB-treated females n=8, 30 months-old MB-treated females n=8. Data tested by multivariate ANOVA presented as mean+/-SEM. Significance accepted at p<0.05 (*p<0.05, **p<0.01, ***p<0.001, ****p<0.0001).

Muscle strength undergoes changes with aging, which can directly or indirectly affect bone. Grip strength is commonly used to measure muscle strength, where the animal holds onto a bar attached to a strain gauge^26^. Previous studies have shown that the administration of 250 µM of MB via drinking water restored the age-related decline in grip strength to levels measured in young mice at 22 months old 19^27, 28^. However, in our study, despite a noticeable decline in grip strength with age, MB did not prevent or improve grip strength (**Figure 6A**).

**Figure 6:**
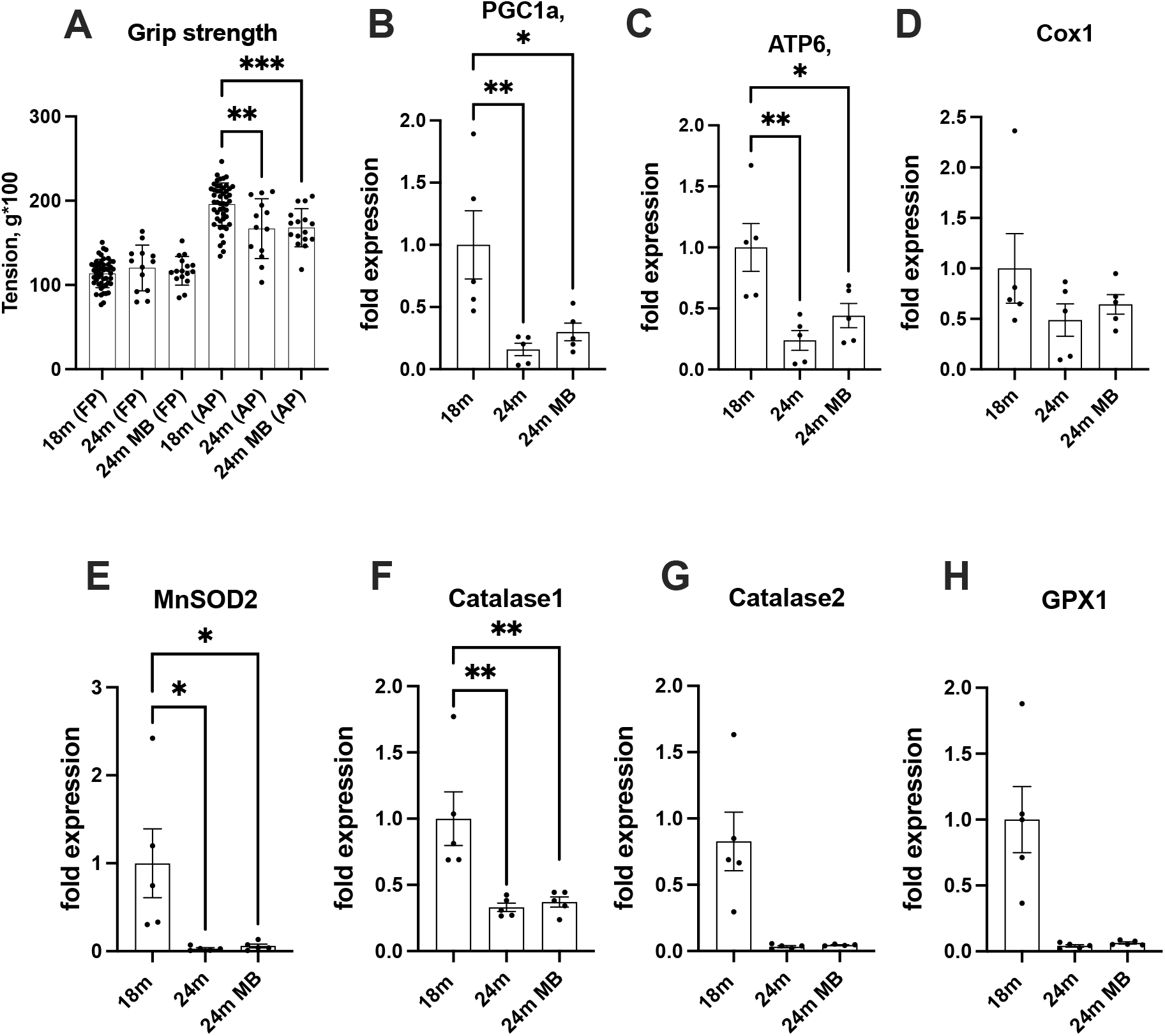
Administration of MB during aging does not affect grip strength or alter skeletal expression of antioxidant enzymes. Eighteen-months old female C57BL/6J were divided into two groups. One group was exposed to regular drinking water and the other to 250 uM MB-containing water for 6 months. (A) Grip strength of the front paws (FP) or all four paws (AP) were determined at 18 (basal) and 24 months using the NBP20-16 Grip Strength apparatus. Data presented as a mean value of three consecutive measurements for FP and AP. 18 months-old females n=48, 24 months-old females n=13, 24 months-old BM-treated females n=16. (B-H) Gene expression was tested in tibia cortical bone by real-time PCR of 18 months old (basal) and 24 months old mice.-including PGC1a-peroxisome proliferator-activated receptor-g coactivator-1a, ATP6-mitochondrial ATP synthase 6, Cox1-cytochrome oxidase subunit 1, MnSOD2-manganase-dependent superoxide dismutase, Catalase 1, Catalase 2, and GPX1-glutathione peroxidase 1. Data tested by multivariate ANOVA and presented as mean+/-SEM of n=5. Significance accepted at p<0.05 (*p<0.05, **p<0.01, ***p<0.01, ****p<0.0001).

Finally, we examined the impact of MB on the expression of genes related to mitochondrial content and function in osteocytes, which are cells embedded within the bone matrix. We observed age-related decreases in the expression of antioxidant enzymes, including catalase-1 and 2, as well as glutathione peroxidase 1 (GPX1). Additionally, the expression of mitochondrial markers, such as ATP synthase 6 (ATP6), cytochrome oxidase subunit 1 (Cox1), peroxisome proliferator-activated receptor-g coactivator-1a (PGC1a), and manganese-dependent superoxide dismutase (MnSOD2), was reduced with age. MB did not have a significant effect on the age-related reductions in the expression of these genes (**Figure 6B-H**).

### Effects of MB and MitoQ on skeletal morphology of the axial and appendicular skeleton in UM-HET3 mice

In order to investigate whether early exposure to MB could lead to improved skeletal outcomes, we collaborated with the Interventions Testing Program (ITP). The ITP is a collaborative effort involving three sites: the University of Michigan (UM) led by Richard A. Miller, the Jackson Laboratory (JL) led by David E. Harrison, and the University of Texas Health Science Center at San Antonio (UT) led by John R Strong. This program aims to identify compounds with translational potential to enhance human health-span during aging.

For our study, we utilized the UM-HET3 mouse model, which is genetically heterogeneous. The UM-HET3 mice are derived from a four-way cross involving CByB6F1 females (JAX stock #100009) and C3D2F1 males (JAX stock #100004) ^29^ (**Figure 1C**). This mouse model has been extensively characterized and documented in various publications (https://phenome.jax.org/projects/ITP1). The advantage of the four-way cross is that each offspring shares 50% of its genome with every other offspring in the population, ensuring population-level diversity and reducing the influence of strain-specific traits on the outcomes.

Using the UM-HET3 mice, we examined the effects of two antioxidant interventions, MB and MitoQ, on skeletal morphology. Male and female UM-HET3 mice were treated with MB (28 ppm administered via food) from 7 to 22 months of age, or MitoQ (100 ppm administered via food) from 4 to 22 months of age, along with their respective control groups. We evaluated the morphology of the axial skeleton (L5) and the appendicular skeleton (femur) using micro-CT (**Table 1**). Long-term administration of MB or MitoQ did not show any significant effects on the skeletal morphology of aged mice. While we did observe significant sex differences in most bone traits at all skeletal sites, there was no significant effect of treatment or interaction between sex and treatment.

**Table 1:**
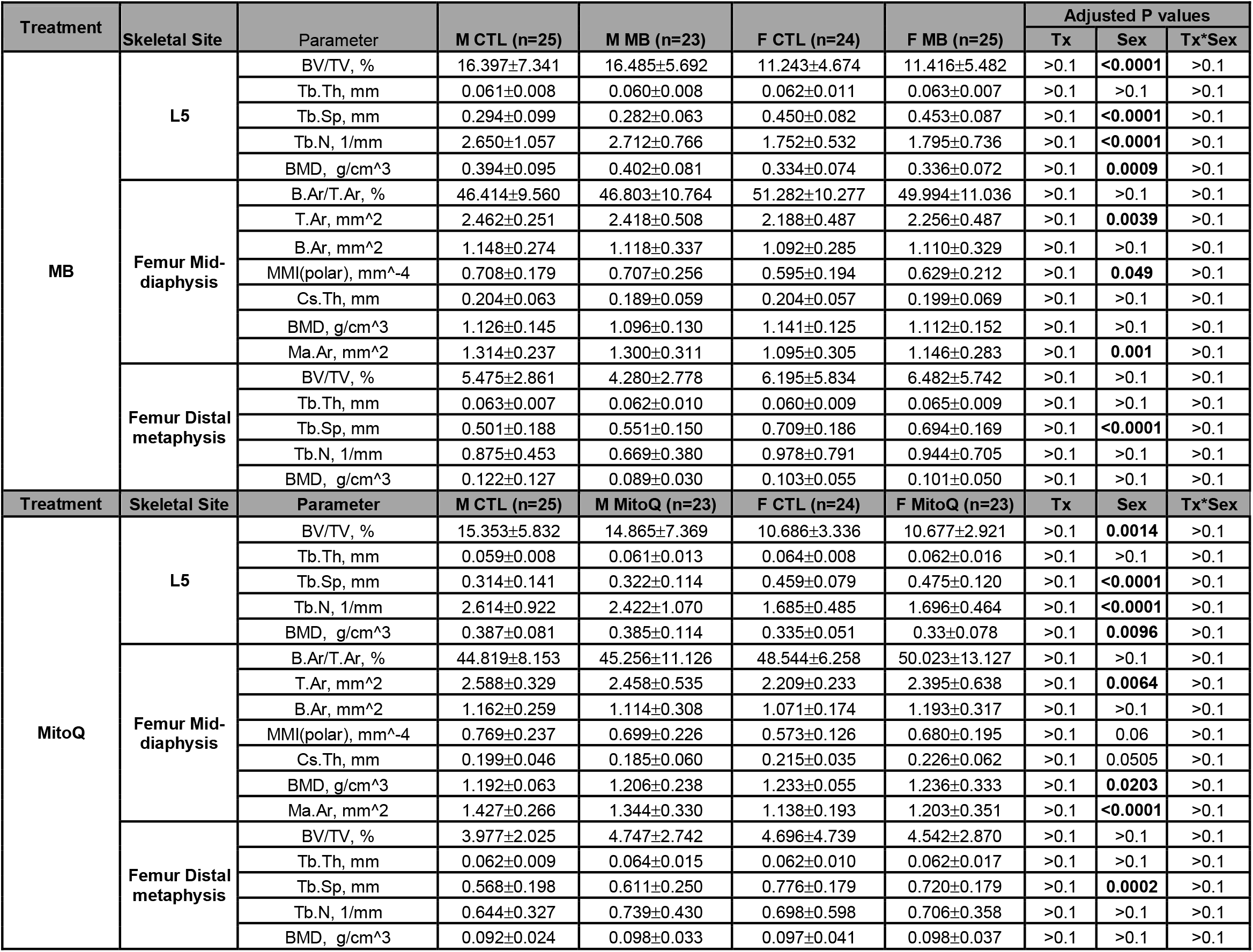
Skeletal morphology of 22-24 months old UM-HET3 mice treated with MB or mitoQ. Femurs and L5 were scanned using SkyScan 1172 (Microphotonics) at 9.7 or 7.5 um resolution, respectively. Data presented as mean±SD of mice from the ITP sites. Data tested by descriptive statistics for this 2×2 factorial design were computed for overall samples, as well as stratified by site and treatment.

## Discussion

The data obtained from the current study suggest that the long-term administration of MB or MitoQ did not have an effect on skeletal morphology during the aging process, even when the treatment was initiated at a young adult age (4 or 7 months-old mice).

Epidemiological studies have provided evidence indicating a relationship between antioxidant intake and bone health. Consequently, numerous studies have been conducted to understand the specific properties of antioxidants and their influence on bone cell metabolism. In vitro studies have demonstrated that exposure to peroxide leads to an increase in bone resorption markers, which can be suppressed by the expression of the antioxidant enzyme catalase^30^. Similarly, overexpression of glutathione peroxidase in osteoclast-like RAW-264.7 cells has been shown to suppress osteoclast differentiation^31^. Peroxide has also been found to inhibit osteoblast differentiation and the formation of type I collagen^32-34^. In vivo studies with C57BL/6J mice have shown a progressive loss of BMD in the spine and femur between 4 and 31 months of age, accompanied by decreased bone remodeling, increased apoptosis of osteoblasts and osteocytes, and heightened oxidative stress^35^. Furthermore, the Forkhead Box O (FoxO) transcription factors, which play a role in defense against oxidative stress, have been found to be crucial for skeletal homeostasis^36^.

MB and MitoQ were tested in UM-HET3 mice by the ITP^9^. Neither agent led to a significant benefit using the log-rank test, but MB did produce a significant increase, in females, in the proportion of mice alive at the 90th percentile, a surrogate for maximum lifespan (links: MB - https://phenome.jax.org/itp/surv/MB/C2009, MitoQ - https://phenome.jax.org/itp/surv/MitoQ/C2015). In many mouse strains, including C57BL/6J and UM-HET3, tumor progression is the primary cause of death, accounting for 70-90% of all mortality cases^37-42^. Although effects of MB and MitoQ on lifespan appear to be limited, at least at the doses tested by the ITP, these drugs might in principle have an effect on age-associated bone loss. A recent study that investigated aging trajectories of various phenotypes and molecular markers throughout the lifespan of male C57BL/6J mice indicated that lifespan data alone cannot be used as an indicator for the effects of specific interventions on age-sensitive phenotypes^43^. This was the rationale behind studying the effects of MB and MitoQ interventions on the skeleton despite their limited influence on lifespan.

Our study had some limitations, including only using a single dose level of MB or MitoQ, as well as only assessing the skeletal phenotype of the UM-HET3 mice once. Additionally, we did not check on the level of oxidative damage in the bone tissue or cells. Nevertheless, using both inbred (C57BL/6J) and genetically heterogeneous (UM-HET3) mouse stocks, we did not observe any evidence that either of these antioxidant strategies had positive impacts on bone mineral density or skeletal morphology during ageing. We are not disputing that redox imbalance within bone cells can be a factor in developing bone disease, but rather that targeting redox balance directly may not be an effective way to prevent age-related bone loss. We think it may be a better course of action to deal with upstream causes of oxidative stress, such as metabolic substrates, which can both yield ROSs and cause impairments in redox homeostasis, when it comes to protecting against age-related decreases in bone health.

## Methods

### Animals

#### Study #1

C57BL/6J female mice: 18 months old C57BL/6J female mice were randomly housed 3-5 in a cage. Mice were housed in a climate-controlled facility with a 12-hour light/dark cycle and provided free access to food and water throughout the experiment. MB (250 µM) was administered via the drinking water. All experiments were approved by The Jackson Laboratory’s IACUC.

#### Study #2

UM-HET3 mice were produced by a cross between (BALB/cByJ × C57BL/6J)F1 mothers (JAX stock #100009) and (C3H/HeJ × DBA/2J)F1 fathers (JAX stock #100004). Detailed housing conditions were specified elsewhere ^44^. All experiments were approved by IACUC of each site.

### Interventions

#### Study #1

Mice were fed irradiated Purina TestDiet 5K0G diet and MB (250 µM, Fluka, through Sigma, St. Louis MO) was administered via the drinking water.

#### Study #2

MB (28 ppm, starting at 4 months of age) or mitoQ (100 ppm, starting at 7 months of age) were administered in the diet. MB and mitoQ were formulated into irradiated Purina TestDiet 5LG6 diet, and supplied to all three sites such that all sites used the same batches of food.

### Micro-CT

Micro-CT was done in accordance with the American Society for Bone and Mineral Research (ASBMR) guidelines ^45^. Bones were scanned using a high-resolution SkyScan micro-CT system (SkyScan 1172, Kontich, Belgium). Femur were scanned at a 9.7μm image voxel size. Cortical bone was analyzed in the mid-diaphysis. Trabecular bone measurements were taken at the femur distal metaphysis 200 μm below the growth plate. The 5^th^ lumbar vertebrae (L5) were scanned at a 7.5 μm image voxel size. Image reconstruction was done using NRecon software (version 1.7.3.0; Bruker micro-CT, Kontich, Belgium), data analysis was done using CTAn software (version 1.17.7.2+; Bruker micro-CT, Kontich, Belgium) and 3D images were constructed using CT Vox software (version 3.3.0 r1403; Bruker micro-CT, Kontich, Belgium).

### Osteoclast cultures

Bone marrow was flushed out of femur and tibia and cultured over-night in alpha-minimum essential medium (α-MEM, Gibco; cat#41061-029)containing 10% fetal bovine serum (FBS, Gibco; cat# 10-438-034), 100 U/mL of penicillin-streptomycin (Gibco; cat# 15-140-122) and 0.25ug/mL amphotericin (Gibco; cat#15-290-026). Nonadherent cells were removed after 24 h in culture, and subjected to a ficoll (GE Healthcare; cat#17-1440-03) density gradient. Mononuclear cells were collected and washed with complete medium and 10^4^ cells per well in 96 wells plate were plated in the presence of 40ng/ml M-CSF(Biolegend; cat#576402) and 60ng/ml RANKL (R&D Systems; cat# 462TEC010) to induce osteoclastogenesis in the presence of different concentrations (0, 0.125 µM, 0.25 µM and 0.5 µM) of MB or MitoQ. TRAP staining (Sigma-Aldrich; cat#387A-1K) was done after 5 days in culture, according to manufacturer’s instructions.

### Bone marrow stromal cell cultures

Bone marrow was flushed out of femur and tibia of 6-7 months old UM-HET3 mice and seeded in α-MEM complete medium. Next day, unattached cells were removed. For a viability assay, cells were collected and seeded in 96 wells plates at the density of 8000 cells per well. Cells were either non-treated (0 µM, Control) or treated with 0.125 µM, 0.25 µM and 0.5 µM MB or MitoQ. At 48 hours, cells were stained with calcein AM (Invitrogen; cat#C3099) and analyzed using spectrophotometer at excitation and emission of 490nm and 520 nm respectively wavelengths. For osteogenesis assays, 0.5 ×10^6^ bone marrow cells were plated on 24 wells plate. After a week, cells were induced for osteogenesis with the supplementation of 50 ug/mL of L-ascorbic acid (Sigma; cat#A5960), 10 nM dexamethasone (Sigma; cat#8893) and 10 mM β-glycro-phosphate in α-MEM complete medium with MB or Mito Q (0, 0.125, 025 and 0.5 uM). The medium was changed twice a week. Alkaline-phosphatase (Alk-Phos) positive cells were assessed on day 18 in culture using Leukocyte Alkaline Phosphatase Kit (Sigma; cat#86R) following manufacturer instructions.

### Mitochondrial stress assay

Primary osteoblast (14 days in culture) were seeded on a type I collagen-coated cell culture micro-plate (4×10^4^ cells/well). Cells were treated with different concentrations (0, 0.125, 0.25 and 0.5 uM) of MB or MitoQ for 48 hours and proceed for mitochondrial stress test (Seahorse Bioscience; cat#101706-100). Accordingly, cells were supplied with 10mM glucose (Sigma; cat#G7528)), 1 mM pyruvate (Sigma; cat#S8636) and 2 mM L-glutamine (Chemicon; cat#TMS-002-C), and oxygen consumption rate (OCR) was determined during basal and upon addition of oligomycin (1 µM), carbonylcyanide p-(trifluoromethoxy) phenylhydrazone (FCCP, 2 µM) and rotenone/antimycin (0.5 µM) using the Seahorse XFe24 bioanalyzer (Agilent). Data from each well was corrected to cell number using the Bio Tek Cytation 1 Cell Imaging Multimode Reader and presented as mean +/-SEM.

**Grip strength** was tested using the NBP20-16 Grip Strength apparatus (Bioseb). Mice were acclimated to the testing room for a minimum of 60 minutes prior to testing. A mouse was gently lowered towards the wire grid and it instinctively grasped the bar with its paws. Care was taken to ensure the mouse is gripping the grid properly, with both front paws only for forepaw measurements or with all four paws for the combined measurements. Once an appropriate grip was assured, the animal was gently and firmly pulled away from the grid by holding the middle of tail until it releases its grasp. Peak force was measured in g during six consecutive trials. Three consecutive forepaw measurements were followed immediately by three consecutive four paw measurements. Average measurements were calculated for each animal. At the conclusion of testing, mice were returned to their cages. The wire grid was cleaned with 70% EtOH after each subject, and sanitized with a CMQ-approved agent (e.g., Virkon) at the end of the test session.

### Gene expression studies

Total RNA was extracted from tibia cortical shells using TRIzol (Invitrogen, Carlsbad, CA, USA) and RNeasy Plus kit (Cat# 74134, Qiagen). cDNA was generated using a commercial kit (Cat# K1621, Thermofisher Scientific). Real-time PCR was done using a BioRad CFX384™ real-time machine with SYBR master mix (Life Technologies/Applied Biosystems, NY, USA Cat#4367659). Transcript levels were assayed triplicates and normalized to *beta-actin*. Primer sequences as follows;

ATP6, Forward: 5’-AGCTCACTTGCCCACTTCCT; Reverse: 5’-AAGCCGGACTGCTAATGCCA-3’, beta-actin, Forward: 5’-GGCTGTATTCCCCTCCATCG-3’; Reverse: 5’-CCAGTTGGTAACAATGCCATGT-3’, Catalase 1, Forward: 5’-GCTGAGAAGCCTAAGAACGC-3’; Reverse:5’-GTCTCCTCAGCGGAGGCTGA-3’, Catalase 2, Forward: 5’-GCAAGTTCCATTACAAGACC-3’; Reverse: 5’-CATAATCCGGATCTTCCTGA-3’, Cox1, Forward: 5’-ATCACTACCAGTGCTAGCCG-3’; Reverse: 5’-CCTCCAGCGGGATCAAAGAA-3’, GPX1, Forward: 5’-GTGGTGCTCGGTTTCCCGTGC-3’; Reverse:5’-CCCGCCACCAGGTCGGACGTA-3’, MnSOD2, Forward: 5’-CTGGCCAAGGGAGATGTTACA-3’ Reverse: 5’-GTCACGCTTGATAGCCTCCAG-3’, and PGC1a, Forward: 5’-TGCAGCGGTCTTAGCACTCA-3’ ; Reverse: 5’-CATGAATTCTCGGTCTTAACAATGG-3’.

### Statistical analyses

Data with inbred (C57BL/6J) mice, presented in figures 2-6 was tested by multivariant ANOVA presented as mean+/-SEM. Significance accepted at p<0.05. Data with genetically heterogeneous (UM-HET3) mice, presented in Table 1. Data presented in table 1 was tested using 2×2 factorial design for overall data as well as stratified by site and treatment. Outcomes included in the study were first analyzed using nonparametric multivariate analysis of variance with permutation tests (10,000 permutations per outcome). Significant multivariate results were subsequently followed up with two-factor aligned ranks transformation analysis of variance with factors for sex, treatment, and the sex*treatment interaction. The false discovery rate for each F-test was controlled for using the method of Benjamini and Hochberg^46^. Finally, p-values from post-hoc tests for each significant factor were adjusted using the Bonferroni method. Analysis was conducted in R v4.1.3.

## Supporting information

Supplementary Figure 1. Effects of MB or MitoQ on BMSCs viability

## Figure legends

Supplementary Figure 1. **Effects of MB or MitoQ on BMSCs viability**. BMSCs isolated from 6-7 months were seeded on 96 wells plate in the presence or absence of MB or Mito Q (0, 0.125, 025, 0.5 uM and 1.0 uM) for 48 hours. Percentage of live cells was determined by calcein AM staining. Data presented as mean+/-SEM of n=3. Data tested by multivariate ANOVA. Significance accepted at p<0.05 (*p<0.05, **p<0.01, ***p<0.001, ****p<0.0001).

## Author contribution

Conceptualization: SY, MBS

Funding acquisition: SY, MBS, RAM, DEH, RLS

Investigation: SBP, DFB, GY, MD

Statistical analyses: RRR

Resources: RK, LR, RAM, DEH, RLS

Writing original draft: SY

Review and editing: All co-authors

## Co-authors details

- Sher Bahadur Poudel

Affiliation: David B. Kriser Dental Center, Department of Molecular Pathobiology, New York University College of Dentistry, New York, NY, 10010, USA

- Dorra Frikha-Benayed

Department of Biomedical Engineering, City College of New York, 160 Convent Avenue, New York, NY, 10031, USA

- Ryan R Ruff

David B. Kriser Dental Center, Department of Epidemiology and Health Promotion, New York University College of Dentistry, New York, NY, 10010, USA

- Gozde Yildirim

Affiliation: David B. Kriser Dental Center, Department of Molecular Pathobiology, New York University College of Dentistry, New York, NY, 10010, USA

- Manisha Dixit

Affiliation: David B. Kriser Dental Center, Department of Molecular Pathobiology, New York University College of Dentistry, New York, NY, 10010, USA

- Mitchell B Schaffler

Department of Biomedical Engineering, City College of New York, 160 Convent Avenue, New York, NY, 10031, USA

- Ron Korstanje

The Jackson Laboratories, Bar Harbor, ME, 04609, USA

- Laura Robinson

The Jackson Laboratories, Bar Harbor, ME, 04609, USA

- Richard A Miller

Pathology and Geriatrics Center, University of Michigan, Ann Arbor, MI, USA.

- David Harrison

The Jackson Laboratory, Bar Harbor, ME, 04609, USA

- Randy Strong

Geriatric Research, Education and Clinical Center and Research Service, South Texas Veterans Health Care System, San Antonio, TX, 78229, USA

Barshop Institute for Longevity and Aging Studies and Department of Pharmacology, The University of Texas Health Science Center, San Antonio, TX, 78229, USA

- Shoshana Yakar

Affiliation: David B. Kriser Dental Center, Department of Molecular Pathobiology, New York University College of Dentistry, New York, NY, 10010, USA

## Notes

**Funding statement:** Financial support received from the National Institutes of Health Grant R01AG056397 to SY and MBS. This work was also supported by the National Institutes of Health grant to The Jackson Laboratory Nathan Shock Center of Excellence in the Basic Biology of Aging AG038070, U01-AG022303 to RAM, UO1-AG022308 to DEH, U01-AG013319 to RS, RS is supported by a Senior Research Career Scientist Award from the Department of Veterans Affairs Office of Research and Development, 1S10 OD026699-01A1 for the Seahorse XFe24 Extracellular Flux Analyzer, and S10 OD010751-01A1 for micro-computed tomography.

**Conflict of interest disclosure:** The authors declare no conflict of interest.

### Competing Interest Statement

The authors have declared no competing interest.

