## Supplementary figures and images for "Targeting mitochondrial dysfunction using methylene blue or mitoquinone to improve skeletal aging"

### Supplementary Figure 1. Effects of MB or MitoQ on BMSCs viability

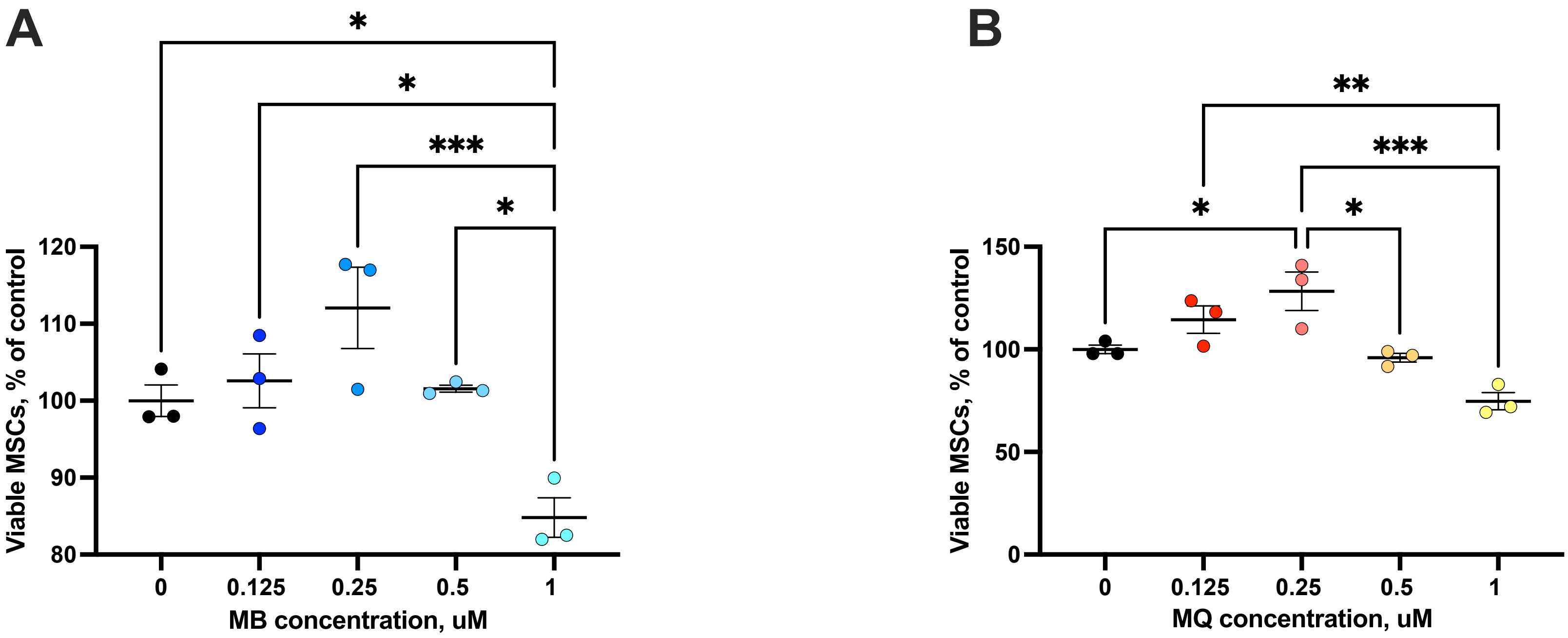
